# Evaluation of *in silico* predictors on short nucleotide variants in *HBA1, HBA2* and *HBB* associated with haemoglobinopathies

**DOI:** 10.1101/2022.04.07.484934

**Authors:** Stella Tamana, Maria Xenophontos, Anna Minaidou, Coralea Stephanou, Cornells L. Harteveld, Celeste Bento, Joanne Traeger-Synodinos, Irene Fylaktou, Norafiza Mohd Yasin, Faidatul Syazlin Abdul Hamid, Ezalia Esa, Hashim Halim-Fikri, Bin Alwi Zilfalil, Andrea C. Kakouri, ClinGen Hemoglobinopathy VCEP, Marina Kleanthous, Petros Kountouris

## Abstract

**Introduction:** Haemoglobinopathies are the commonest monogenic diseases worldwide and are caused by variants in the globin gene clusters. With over 2400 variants detected to date, their interpretation using the ACMG/AMP guidelines is challenging, with computational evidence able to provide valuable input about their functional annotation. While many *in silico* predictors have already been developed, their performance varies for different genes and diseases.

**Materials and Methods:** We evaluate 31 *in silico* predictors using a dataset of 1627 variants in *HBA1, HBA2,* and *HBB.* Through varying the decision threshold for each tool, we analyse their performance (a) as binary classifiers of pathogenicity, and (b) using different nonoverlapping pathogenic and benign thresholds for their optimal use in the ACMG/AMP framework.

**Results:** CADD, Eigen-PC, and REVEL are the overall top performers, with the former reaching moderate strength level for pathogenic prediction. Eigen-PC and REVEL achieve the highest accuracies for missense variants, while CADD is also a reliable predictor of non-missense variants. Moreover, SpliceAI is the top performing splicing predictor, reaching strong level of evidence, while GERP++ and phyloP are the most accurate conservation tools.

**Discussion:** This study provides evidence about the optimal use of computational evidence in globin gene clusters under the ACMG/AMP framework.

## Introduction

With genetic testing frequently employed by clinical laboratories to aid diagnosis and treatment decisions in different diseases^1^, advances in sequencing technology produce an excessive amount of sequencing data leading to a rapidly enlarging pool of new unclassified variants. While sequencing data provide new candidates for therapeutic interventions and personalized medicine, they also introduce challenges in correctly classifying variants as pathogenic or benign. Thus, variant interpretation often relies on human expertise to gather information from different and diverse sources as to combine individual pieces of evidence into a comprehensive estimate with high confidence^2^.

To assist in the establishment of a common framework for standardized variant classification, the American College of Medical Genetics and Genomics (ACMG) and the Association for Molecular Pathology (AMP) published joint recommendations for the interpretation of genetic variants^1^. The ACMG/AMP framework was designed for use across different genes and diseases, thus requiring further specification in disease-specific scenarios. In response to this need, the Clinical Genome (ClinGen) Resource formed various disease-specific variant curation expert panels (VCEPs) to develop specifications to the ACMG/AMP framework^3^. The ClinGen Hemoglobinopathy VCEP focuses on performing and testing the applicability of haemoglobinopathy-specific modifications to the standard ACMG/AMP framework before proceeding with the classification and interpretation of variants related to haemoglobinopathies^4^. Haemoglobinopathies represent the commonest groups of inherited monogenic disorders affecting approximately 7% of the global population^5^. They are caused by genetic defects in genes located in the α-globin locus (Accession: NG_000006) and in the β-globin locus (Accession: NG_000007). To date, there are over 2400 different naturally occurring globin gene variants, which are collected and manually curated in IthaGenes, a haemoglobinopathy-specific database on the ITHANET portal^6^.

The ACMG/AMP guidelines propose the use of *in silico* predictors (namely criteria PP3 and BP4 for pathogenic and benign evidence, respectively) as supporting evidence for variant pathogenicity classification^1^. Several tools have already been developed to predict the impact of genetic variants and their relation to developing diseases. These tools fall into four main categories based on the theoretical background and type of data they use for predicting variant effect, namely sequence conservation-based, structure-based analysis, combined (i.e., including both sequence and structural features), and meta-predictors^7^.

The performance of different *in silico* tools varies across genes and diseases as numerous studies illustrated discrepancies regarding variant pathogenicity prediction^2,8–11^. Previous studies have also evaluated the performance of *in silico* predictors for globin gene variants^12,13^, demonstrating a high degree of discordance between *in silico* tools. Therefore, it is evident that a disease- or gene-specific evaluation of *in silico* tools can provide evidence for the optimal selection or combination of tools to identify the functional impact of variants. Recently, ClinGen published a study on the performance of four in *silico* predictors using a set of 237 variants^14^, suggesting that custom thresholds should be explored for each *in silico* tool to establish PP3 and BP4 criteria. However, given the impact of *in silico* tools on variant classification, further calibration with larger datasets is still needed to optimize their performance.

The main purpose of this study is to compare the performance of various *in silico* predictors and determine the most appropriate ones for predicting the functional impact of SNVs in *HBA1, HBA2,* and *HBB* related to haemoglobinopathies. We selected 31 *in silico* predictors, including those recommended by ClinGen^3^ and linked in the Variant Curation Interface (VCI)^15^, along with additional tools described in literature. A total of 1627 short nucleotide variants (SNVs) were retrieved from the IthaGenes database^6,16^ and were annotated using a Delphi approach with respect to their pathogenicity by experts (co-authoring this study) involved in haemoglobinopathy molecular diagnosis in five different countries. The annotated pathogenicity of each SNV was then used to evaluate its predicted pathogenicity provided by *in silico* tools. To our knowledge, this is the largest comparative study of *in silico* tools for SNVs related to haemoglobinopathies in terms of both the number of tools used and the size of utilised variant dataset.

## Methods

### Dataset

Figure 1 shows a schematic representation of the main steps of our methodology. SNVs were retrieved from the IthaGenes database of the ITHANET portal^6,16^. The dataset includes all SNVs (≤50bp) curated in IthaGenes (access date: 05/02/2021) located in *HBA1, HBA2,* and *HBB,* excluding (i) disease-modifying variants, (ii) complex variants with multiple DNA changes found *in cis,* and (iii) variants whose genomic location is unclear, such as α-chain variants identified by protein studies without identifying the affected α-globin gene.

**Figure 1.**
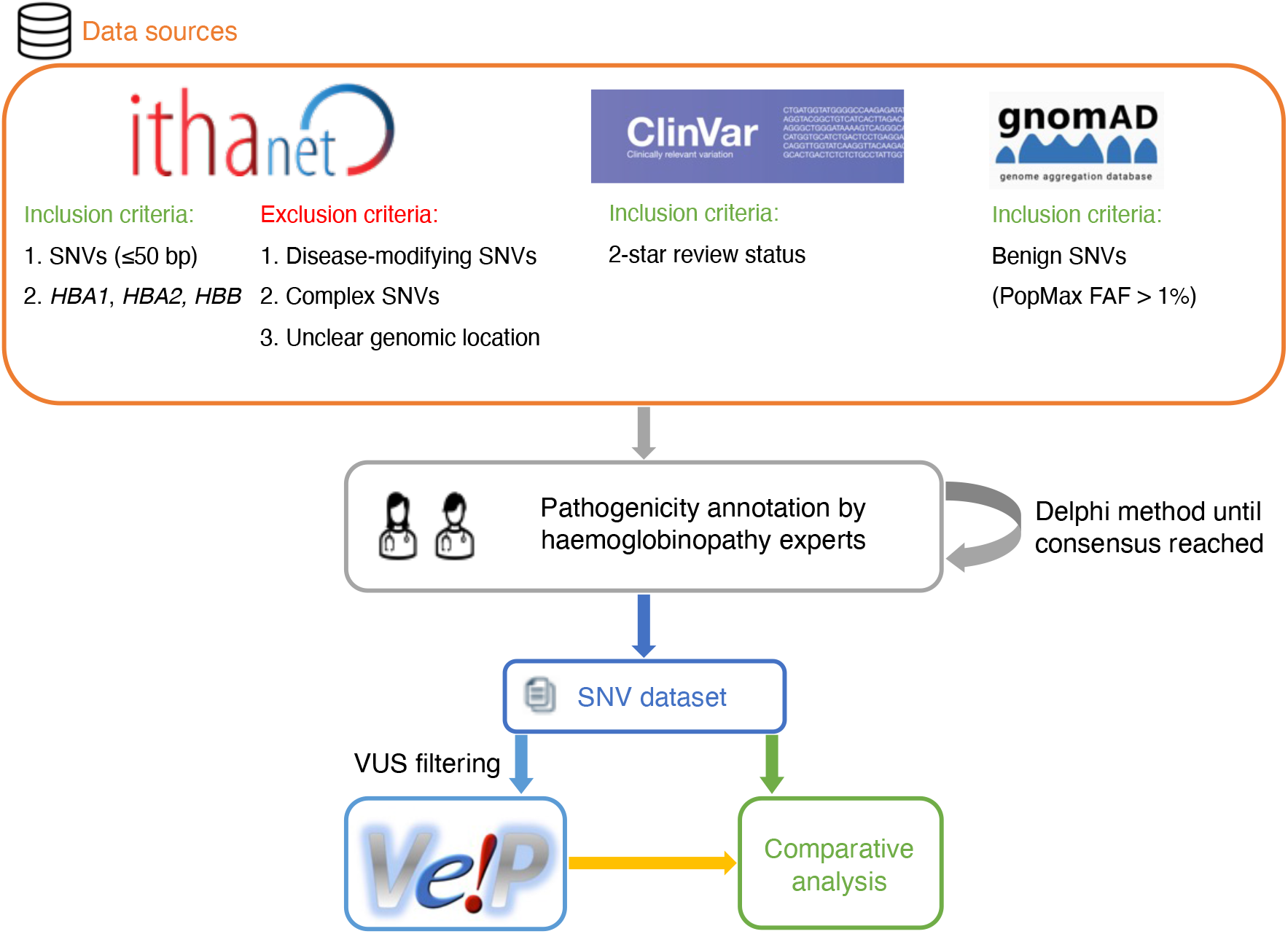
A schematic flowchart of the methodology followed for this comparative analysis.

Additionally, we queried ClinVar (access date: 05/02/2021)^17^ for SNVs with a two-star review status and gnomAD (access date: 05/02/2021)^18^ for benign/likely benign SNVs using PopMax Filtering Allele Frequency (FAF) greater than 1% in *HBA1, HBA2* and *HBB.* Any missing SNVs were added to both IthaGenes and the dataset of this study. The final dataset included 1627 distinct SNVs. Finally, the dataset was further processed using the batch service of Variant Validator^19^ to validate the HGVS names and correct any annotation errors.

### Annotated variant pathogenicity

To enable the evaluation of *in silico* predictions, we subsequently annotated the pathogenicity of each SNV and compared it to the results of *in silico* predictors. Specifically, we used existing curated information on IthaGenes and further collected available evidence in scientific literature for each SNV in the dataset. The pathogenicity for each SNV was annotated using the following criteria:

- **Pathogenic/Likely pathogenic (P/LP)**

○ SNVs that result in abnormal haematology or abnormal Hb properties, or sometimes causing disease (i.e., dominant), when detected in heterozygotes, OR
○ Causes disease when observed *in trans* with an established pathogenic variant or in the homozygous state
- **Benign/Likely Benign (B/LB)**

○ At least three (independent) occurrences of the variant in heterozygous state without any change in the haematological parameters and Hb properties OR
○ Not causing disease when observed *in trans* with an established pathogenic variant
- **Variant of uncertain significance (VUS)**

○ All variants that do not meet the above criteria for benign/pathogenic or have conflicting evidence

Importantly, we used a Delphi approach^20^ to allow independent evaluation of the curated evidence for each variant. The pathogenicity of each SNV was independently assessed by two different groups of haemoglobinopathy experts, using evidence curated by the IthaGenes database or collected as part of this study. Then, the independent annotations were merged into one consensus classification. In cases of disagreement, a consensus pathogenicity status was decided, after discussion among all experts, or the SNV was marked as a VUS. SNVs that have been directly submitted to IthaGenes by experts not participating in this study and without a peer-reviewed publication describing the methodology and results, have been also annotated as VUS. After descriptive analysis of the full dataset, 601 SNVs annotated as VUS were filtered out of the dataset.

For the evaluation of tools predicting the impact of variants on splicing, we further annotated variants with respect to their effect on gene/protein function and assembled the following datasets:

a. Variants affecting splicing: all P/LP variants annotated to affect splicing or being in the splicing region of the transcript, excluding variants that are annotated as both missense and splicing and, therefore the mechanism of pathogenicity is ambiguous
b. Variants not affecting splicing: all remaining variants in the dataset (P/LP and B/LB), excluding those annotated as both missense and splicing.

For SpliceAI, we selected the highest of the four Delta Scores (DS) provided as output, while for MaxEntScan we used two different thresholds as follows: (a) the absolute difference between the reference and alternative allele (denoted as Diff), and (b) the absolute percentage of change between the reference and alternative allele (denoted as Per)^21^.

### *In silico* prediction tools

Thirty one *in silico* predictors were compared in this study, as follows: ada^22^, BayesDel^23^, CADD^24^, ClinPred^25^, CONDEL^26^, DANN^27^, EIGEN-PC^28^, FATHMM^29^, FATHMM-MKL^30^, fitCons^31^, GERP++^32^, LIST-S2^33^, LRT^34^, MaxEntScan^35^, Meta-SVM^36^, MetaLR^37^, MutationAssessor^38^, MutationTaster^39^, MutPred2^40^, PolyPhen-2^41^, PROVEAN^42^, REVEL^43^, rf^22^, SIFT^44^, SpliceAI^45^, VEST4^46^, phastCons (phastCons17way and phastCons30way)^47^, phyloP (phyloP100way and phyloP30way)^47^, and SiPhy_29way^48^. Four of the tools are focused on predicting the splicing impact of a variant (ada, MaxEntScan, rf, and SpliceAI), while six tools produce conservation scores (GERP++, phastCons17way, phastCons30way, phyloP100way, phyloP30way, and SiPhy_29way). We selected *in silico* tools recommended by ClinGen and available in the ClinGen VCI^15^, as well as additional established tools used in previous studies. We employed the online version of the Ensembl VEP^49^ and its dbNSFP^50^ plugin (version 4.2a) to obtain the prediction scores of the variants in our dataset.

### Predictive performance assessment

Commonly used scalar measures were employed to compare the prediction accuracy of *in silico* tools, including specificity, sensitivity, and accuracy. All of them can be derived from two or more of the following quantities: (1) true positives (TP), the number of correctly predicted P/LP variants; (2) true negatives (TN), the number of correctly predicted B/LB variants; (3) false positives (FP), the number of B/LB variants incorrectly predicted as P/LP; (4) false negatives (FN), the number of P/LP variants incorrectly predicted as B/LB. Specificity is defined as the fraction of correctly predicted B/LB variants, sensitivity is the fraction of correctly predicted P/LP variants, and accuracy is the ratio of correct predictions versus the total number of predictions^51^.

Moreover, we used the Matthews Correlation Coefficient (MCC)^52^ to compare the performance of *in silico* predictors. MCC ranges from −1 (i.e., always falsely predicted) to 1 (i.e., perfectly predicted) with a value of 0 corresponding to random prediction. MCC is considered one of the most robust measures to evaluate binary classifiers^53^. Hence, in our analysis, the optimal threshold for binary classification was the one that maximised the MCC for each *in silico* tool.

Following the guidelines of a Bayesian variant classification framework^54^, likelihood ratios (LRs) for pathogenic (LR+) and benign (LR-) outcomes were calculated for each tool to evaluate the evidence strength of their pathogenicity prediction using the Odds of Pathogenicity (OddsP) in the Bayesian framework. According to the Bayesian framework, the strength of OddsP for each evidence level was set as follows: ‘Very Strong’ (350:1), ‘Strong’ (18.7:1), ‘Moderate’ (4.3:1) and ‘Supporting’ (2.08:1).

### Comparative Analysis

The analysis was separated into three parts. First, we performed descriptive analysis of the dataset, including variants annotated as VUS, based on the variant type, the variant effect on gene/protein function, the haemoglobinopathy disease group, thalassemia phenotype, molecular mechanism, and annotated pathogenicity. Subsequently, we removed variants annotated as VUS and we compared the 31 *in silico* tools as binary predictors of variant pathogenicity by selecting the threshold that maximized the MCC for each tool. For predictors whose output scores ranged from 0 to 1, we used thresholds with intervals of 0.05, whereas for predictors with scores falling outside this range, we set 19 custom ranges based on the observed minimum and maximum scores in our dataset. Finally, we identified separate non-overlapping thresholds for prediction of pathogenic and benign effect as recommended by the Bayesian framework for variant interpretation^54^, by selecting thresholds passing the recommended LR+ and LR- thresholds, while maximising the percentage of correctly predicted variants for each tool. For tools passing the LR thresholds, we further finetuned the decision thresholds using smaller steps to optimise the prediction accuracy. Statistical analysis and visualisation of the results were performed using custom R scripts and the epiR package.

## Results

### Descriptive analysis

Initially, we performed a descriptive analysis of the full dataset, including variants annotated as VUS, which comprised 1627 SNVs. In terms of the annotated pathogenicity, 194 (11.9%) SNVs classified as B/LB, 832 (51.1%) as P/LP and 601 (36.9%) as VUS. The distribution per globin gene is the following: 553 P/LP, 77 B/LB, and 403 VUS for *HBB* (total: 1033 SNVs; 63.5%), 173 P/LP, 66 B/LB, and 111 VUS for *HBA2* (total: 350 SNVs; 21.5%), and 106 P/LP, 51 B/LB, and 87 VUS for *HBA1* (total: 245 SNVs; 15%). Supplementary Figure 1 illustrates the distribution of variants on each globin gene based on their annotated pathogenicity and demonstrates the highest fraction of P/LP variant in protein coding regions and in canonical splice sites. Increased numbers of P/LP variants are also observed in specific noncoding regions of the globin genes, such as polyadenylation regions and the promoter and 5’ UTR for *HBB.*

Figure 2 summarises the distribution of SNVs in the dataset according to their effect on gene/protein function with respect to the annotated pathogenicity (Panel A), the annotated haemoglobinopathy group (Panel B), the thalassaemia allele phenotype (Panel C), altered oxygen affinity (Panel D), altered stability (Panel E), and the molecular mechanism involved in pathogenesis (Panel F). The effect on gene/protein function includes the following categories: (a) missense variants (SO:0001583), (b) synonymous variants (SO:0001819), (c) frameshift (SO:0001589), (d) initiation codon (SO:0000318), (e) in-frame indels (SO:0001820), (f) splicing, including cryptic splice site (SO:0001569), splice acceptor (SO:0001574), splice donor (SO:0001575) and splice region variants (SO:0001630), (g) stop lost (SO:0001578), (h) stop gained (SO:0001587), and (i) variants in regulatory elements, including promoter (SO:0001631), 5’ UTR (SO:0001623), 3’ UTR (SO:0001624) and polyadenylation variants (SO:0001545). Importantly, there are no B/LB null variants (i.e., frameshifts, stop gained, canonical splice sites, initiation codon) in the dataset, which reflects that loss-of-function is a primary disease mechanism, particularly for thalassaemia syndromes. In contrast, missense variants, representing the largest variant type category (total: 960 SNVs; 59%), are present in all pathogenicity categories, with 115 (12% of SNVs in the category), 331 (34.5%), and 514 (53.5%) annotated as B/LB, P/LP, and VUS, respectively. The distribution of missense variants in the three categories and the high percentage of missense VUS highlight the challenge to interpret the pathogenicity of missense variants in the globin genes, requiring rigorous study of available evidence, including computational evidence.

**Figure 2.**
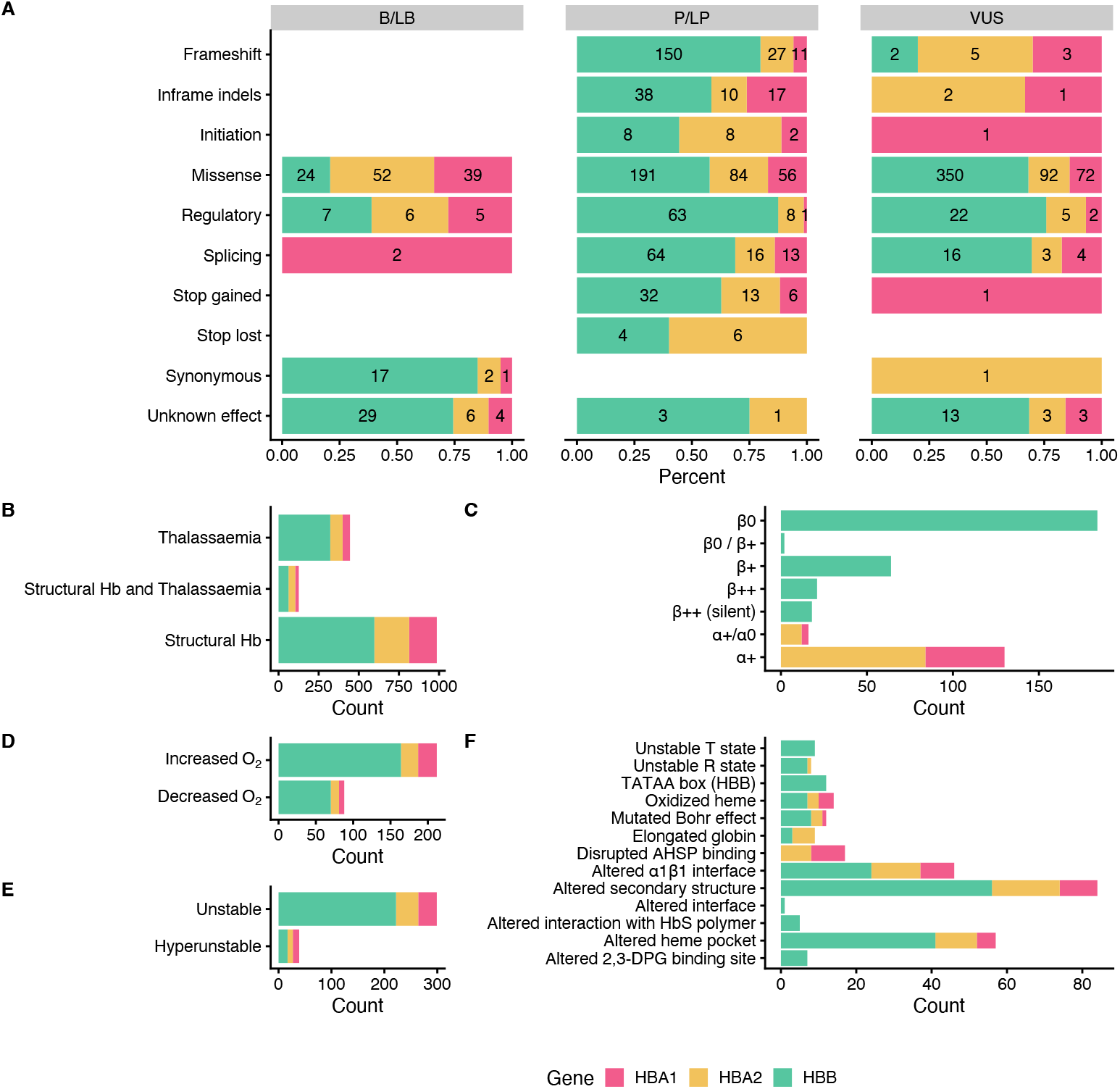
Descriptive plots of the SNV dataset. (A) Variant effect on gene/protein function with respect to the annotated pathogenicity status. (B) Haemoglobinopathy group, (C) Thalassaemia phenotype, (D) O2 affinity (E) Hb stability, (F) Molecular mechanisms

Moreover, the dataset comprises SNVs causing structural haemoglobinopathies (986 SNVs), thalassaemia (445 SNVs), and both thalassaemia and structural haemoglobinopathies (128 SNVs). The thalassaemia phenotype group describes the allele phenotype and includes *HBA1* and *HBA2* variants (α^+^/α° and α^+^; total: 146 SNVs) and *HBB* variants (β°, β°/β^+^, β+, β++ (silent) and β++; total: 289 SNVs). Here, we observed that most variants have allele phenotype of α^+^ (130 SNVs) and β° (184 SNVs). The category of Hb stability is further divided into hyperunstable (39 SNVs) and unstable (299 SNVs), while the Hb O_2_ affinity group is divided into increased O_2_ affinity (212 SNVs) and decreased O_2_ affinity (88 SNVs). The main molecular mechanisms disrupted are alterations of the secondary structure (84 SNVs), heme pocket (57 SNVs), and α1β1 interface (46 SNVs). The disruption of the molecular mechanisms has been associated with clinical phenotypes, such as haemolytic anaemia, reticulocytosis, erythrocytosis, and cyanosis^55^.

### Evaluation of *in silico* tools as binary predictors

Table 1 shows a comparison of all *in silico* predictors used in this study as binary classifiers of pathogenicity, against the consensus dataset with VUS removed. For each tool, we varied the decision threshold for the whole range of possible prediction scores and calculated all statistical measures in each step (Supplementary Table 2). For binary pathogenicity classification, we selected the threshold that maximised the MCC for each tool. Accuracy ranged from 51% (FATHMM) to 84% (CADD) with a median value of 76%. The sensitivity ranged from 41% (FATHMM) to 100% (fitCons) with a median of 82.5%, while specificity ranged from 1% (fitCons) to 81% (BayesDel) with a median of 54%. High sensitivity and low specificity indicate that most predictors correctly predict the P/LP variants, but misclassify the B/LB ones. MCC values ranged from 0.04 (fitCons), indicating almost random prediction, to 0.49 (CADD) with a median value of 0.32. CADD achieved the highest accuracy and MCC among all *in silico* tools tested, using the threshold maximising the MCC (>10.44 for pathogenic prediction), indicating good performance as a binary classifier for globin gene variants. However, this threshold is not optimal for predicting benign variants, with the achieved specificity (0.47) being below the median, hence misclassifying 101 out of 192 B/LB SNVs. Eigen-PC achieved the second highest MCC (0.44), sensitivity of 0.79 and specificity of 0.7, with decision threshold of 1.87.

**Table 1.**
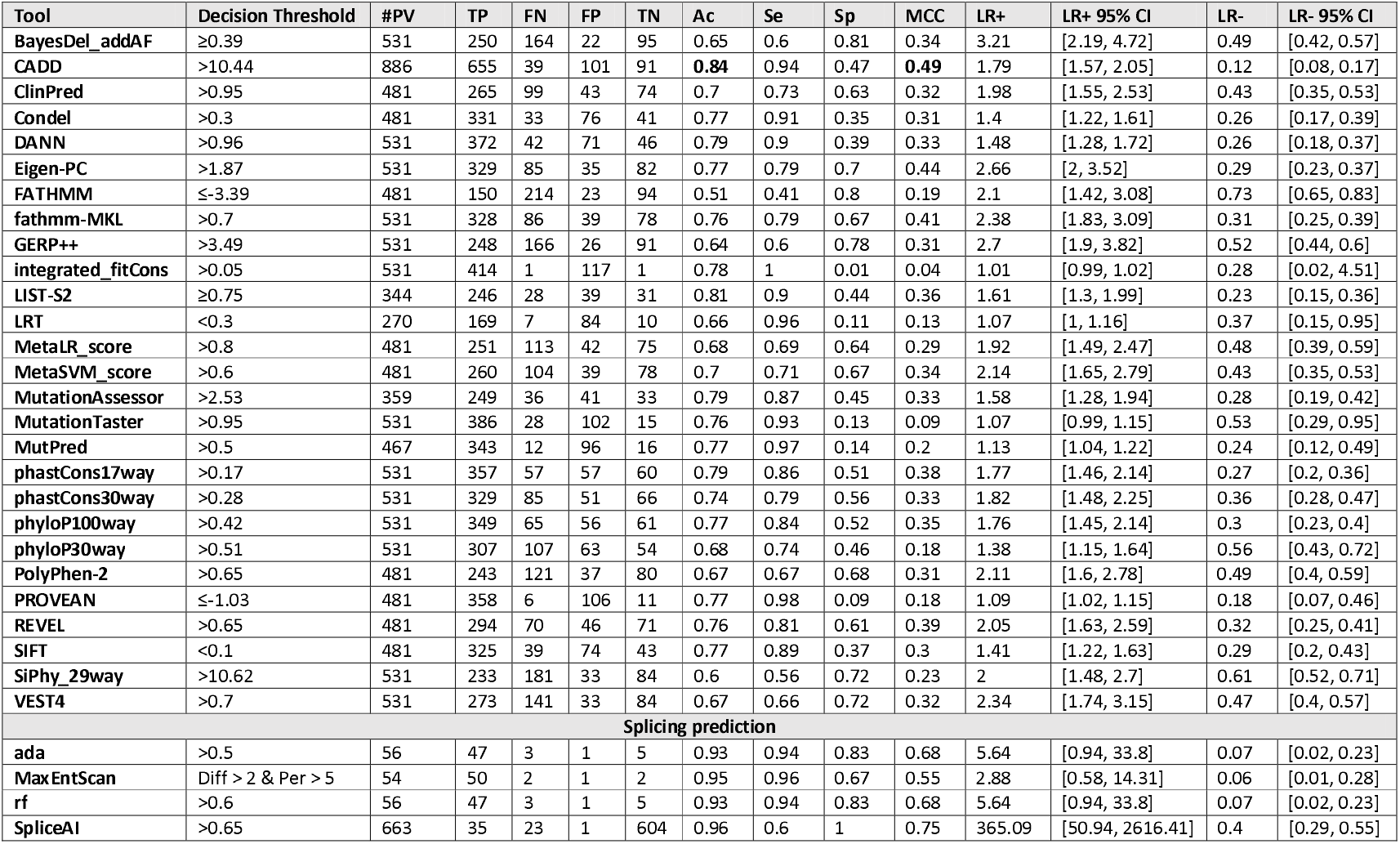
Results and performance comparison of *in silico* predictors with the optimal threshold based on MCC. **#PV:** number of predicted variants; **Ac:** Accuracy; **Se:** Sensitivity; **Sp:** Specificity; **MCC:** Matthews Correlation Coefficient; **LR^+^**: positive Likelihood Ratio; **LR-:** negative Likelihood Ratio; **95% CI:** 95% Confidence Interval.

| Tool | Decision Threshold | #PV | TP | FN | FP | TN | Ac | Se | Sp | MCC | LR+ | LR+ 95% CI | LR- | LR- 95% CI |
| --- | --- | --- | --- | --- | --- | --- | --- | --- | --- | --- | --- | --- | --- | --- |
| BayesDel_addAF | ≥0.39 | 531 | 250 | 164 | 22 | 95 | 0.65 | 0.6 | 0.81 | 0.34 | 3.21 | [2.19, 4.72] | 0.49 | [0.42, 0.57] |
| CADD | >10.44 | 886 | 655 | 39 | 101 | 91 | **0.84** | 0.94 | 0.47 | **0.49** | 1.79 | [1.57, 2.05] | 0.12 | [0.08, 0.17] |
| ClinPred | >0.95 | 481 | 265 | 99 | 43 | 74 | 0.7 | 0.73 | 0.63 | 0.32 | 1.98 | [1.55, 2.53] | 0.43 | [0.35, 0.53] |
| Condel | >0.3 | 481 | 331 | 33 | 76 | 41 | 0.77 | 0.91 | 0.35 | 0.31 | 1.4 | [1.22, 1.61] | 0.26 | [0.17, 0.39] |
| DANN | >0.96 | 531 | 372 | 42 | 71 | 46 | 0.79 | 0.9 | 0.39 | 0.33 | 1.48 | [1.28, 1.72] | 0.26 | [0.18, 0.37] |
| Eigen-PC | >1.87 | 531 | 329 | 85 | 35 | 82 | 0.77 | 0.79 | 0.7 | 0.44 | 2.66 | [2, 3.52] | 0.29 | [0.23, 0.37] |
| FATHMM | ≤-3.39 | 481 | 150 | 214 | 23 | 94 | 0.51 | 0.41 | 0.8 | 0.19 | 2.1 | [1.42, 3.08] | 0.73 | [0.65, 0.83] |
| fathmm-MKL | >0.7 | 531 | 328 | 86 | 39 | 78 | 0.76 | 0.79 | 0.67 | 0.41 | 2.38 | [1.83, 3.09] | 0.31 | [0.25, 0.39] |
| GERP++ | >3.49 | 531 | 248 | 166 | 26 | 91 | 0.64 | 0.6 | 0.78 | 0.31 | 2.7 | [1.9, 3.82] | 0.52 | [0.44, 0.6] |
| integrated_fitCons | >0.05 | 531 | 414 | 1 | 117 | 1 | 0.78 | 1 | 0.01 | 0.04 | 1.01 | [0.99, 1.02] | 0.28 | [0.02, 4.51] |
| LIST-S2 | ≥0.75 | 344 | 246 | 28 | 39 | 31 | 0.81 | 0.9 | 0.44 | 0.36 | 1.61 | [1.3, 1.99] | 0.23 | [0.15, 0.36] |
| LRT | <0.3 | 270 | 169 | 7 | 84 | 10 | 0.66 | 0.96 | 0.11 | 0.13 | 1.07 | [1, 1.16] | 0.37 | [0.15, 0.95] |
| MetaLR_score | >0.8 | 481 | 251 | 113 | 42 | 75 | 0.68 | 0.69 | 0.64 | 0.29 | 1.92 | [1.49, 2.47] | 0.48 | [0.39, 0.59] |
| MetaSVM_score | >0.6 | 481 | 260 | 104 | 39 | 78 | 0.7 | 0.71 | 0.67 | 0.34 | 2.14 | [1.65, 2.79] | 0.43 | [0.35, 0.53] |
| MutationAssessor | >2.53 | 359 | 249 | 36 | 41 | 33 | 0.79 | 0.87 | 0.45 | 0.33 | 1.58 | [1.28, 1.94] | 0.28 | [0.19, 0.42] |
| MutationTaster | >0.95 | 531 | 386 | 28 | 102 | 15 | 0.76 | 0.93 | 0.13 | 0.09 | 1.07 | [0.99, 1.15] | 0.53 | [0.29, 0.95] |
| MutPred | >0.5 | 467 | 343 | 12 | 96 | 16 | 0.77 | 0.97 | 0.14 | 0.2 | 1.13 | [1.04, 1.22] | 0.24 | [0.12, 0.49] |
| phastCons17way | >0.17 | 531 | 357 | 57 | 57 | 60 | 0.79 | 0.86 | 0.51 | 0.38 | 1.77 | [1.46, 2.14] | 0.27 | [0.2, 0.36] |
| phastCons30way | >0.28 | 531 | 329 | 85 | 51 | 66 | 0.74 | 0.79 | 0.56 | 0.33 | 1.82 | [1.48, 2.25] | 0.36 | [0.28, 0.47] |
| phyloP100way | >0.42 | 531 | 349 | 65 | 56 | 61 | 0.77 | 0.84 | 0.52 | 0.35 | 1.76 | [1.45, 2.14] | 0.3 | [0.23, 0.4] |
| phyloP30way | >0.51 | 531 | 307 | 107 | 63 | 54 | 0.68 | 0.74 | 0.46 | 0.18 | 1.38 | [1.15, 1.64] | 0.56 | [0.43, 0.72] |
| PolyPhen-2 | >0.65 | 481 | 243 | 121 | 37 | 80 | 0.67 | 0.67 | 0.68 | 0.31 | 2.11 | [1.6, 2.78] | 0.49 | [0.4, 0.59] |
| PROVEAN | ≤-1.03 | 481 | 358 | 6 | 106 | 11 | 0.77 | 0.98 | 0.09 | 0.18 | 1.09 | [1.02, 1.15] | 0.18 | [0.07, 0.46] |
| REVEL | >0.65 | 481 | 294 | 70 | 46 | 71 | 0.76 | 0.81 | 0.61 | 0.39 | 2.05 | [1.63, 2.59] | 0.32 | [0.25, 0.41] |
| SIFT | <0.1 | 481 | 325 | 39 | 74 | 43 | 0.77 | 0.89 | 0.37 | 0.3 | 1.41 | [1.22, 1.63] | 0.29 | [0.2, 0.43] |
| SiPhy_29way | >10.62 | 531 | 233 | 181 | 33 | 84 | 0.6 | 0.56 | 0.72 | 0.23 | 2 | [1.48, 2.7] | 0.61 | [0.52, 0.71] |
| VEST4 | >0.7 | 531 | 273 | 141 | 33 | 84 | 0.67 | 0.66 | 0.72 | 0.32 | 2.34 | [1.74, 3.15] | 0.47 | [0.4, 0.57] |
| **Splicing prediction** |  |  |  |  |  |  |  |  |  |  |  |  |  |  |
| ada | >0.5 | 56 | 47 | 3 | 1 | 5 | 0.93 | 0.94 | 0.83 | 0.68 | 5.64 | [0.94, 33.8] | 0.07 | [0.02, 0.23] |
| MaxEntScan | Diff > 2 & Per > 5 | 54 | 50 | 2 | 1 | 2 | 0.95 | 0.96 | 0.67 | 0.55 | 2.88 | [0.58, 14.31] | 0.06 | [0.01, 0.28] |
| rf | >0.6 | 56 | 47 | 3 | 1 | 5 | 0.93 | 0.94 | 0.83 | 0.68 | 5.64 | [0.94, 33.8] | 0.07 | [0.02, 0.23] |
| SpliceAI | >0.65 | 663 | 35 | 23 | 1 | 604 | 0.96 | 0.6 | 1 | 0.75 | 365.09 | [50.94, 2616.41] | 0.4 | [0.29, 0.55] |

When used as binary predictors, the *in silico* tools were unable to reach the strength level required by the Bayesian Framework^54^ to provide supporting evidence for variant classification. Although four tools (Eigen-PC, fathmm-MKL, VEST4, MetaSVM) achieved LR+ higher than 2.08 and LR- lower than 0.48, required for supporting evidence strength for pathogenic and benign classification, respectively, their 95% Confidence Intervals (95% Cl) extended beyond the above thresholds and, therefore, are not recommended alone for variant interpretation.

Supplementary Figure 2 shows a heatmap illustrating the extend of concordance among 27 *in silico* tools (excluding splicing tools) and clustering of the tools based on their concordance, using the thresholds that maximised the MCC (Table 1). Notably, we observe a high degree of concordance for P/LP variants in *HBB* (top of the heatmap), while there is a lower degree of concordance for variants in *HBA1* and *HBA2* (middle of the heatmap). The bottom part of the heatmap illustrates a higher discordance for B/LB variants in *HBA1* and *HBA2.*

### Performance of splicing predictors

Table 1 summarises the performance of *in silico* splicing tools using the threshold that maximised the MCC. With most SNVs affecting splicing regions of the globin genes annotated as P/LP, the performance of splicing tools cannot be compared reliably because of the limited number of negative examples in the dataset, i.e. B/LB SVNs in splicing regions. Out of the four *in silico* tools tested, only SpliceAI provides a prediction score for variants that are not located near the canonical splicing sites. All splicing effect predictors displayed high accuracy, ranging from 93% (ada and rf) to 96% (SpliceAI), moderate to high sensitivity, ranging from 0.6 (SpliceAI) to 0.96 (MaxEntScan), and moderate to high specificity ranging from 0.67 (MaxEntScan) to 1 (SpliceAI). The MCC values ranged from 0.55 (MaxEntScan) to 0.75 (SpliceAl). SpliceAl achieved a high LR+ indicating strong performance in predicting SNVs disrupting splicing. The low number (≤5) of TN, FP and FN in the predictions make the calculation of LRs for the remaining tools unreliable.

### Evaluation with different pathogenic and benign thresholds

We subsequently calibrated separate non-overlapping thresholds for pathogenic and benign prediction for each *in silico* tool to maximise both the percentage of variants correctly predicted by the selected threshold pairs that meet at least the supporting strength LR thresholds as defined by the Bayesian framework. More specifically, we filtered tools that achieved a lower bound 95% CI LR+ of 2.08 or higher for pathogenic prediction and an upper bound 95% CI LR- of 0.48 or lower for benign prediction. Figure 3A illustrates the changing LR values for the nine tools that reached these thresholds, while varying the decision thresholds. For these tools, we further finetuned the decision thresholds using smaller steps for the varying thresholds to maximise the number of correctly predicted SNVs. Furthermore, we tested the performance of all tools in different subsets of the dataset, including missense-only, non-missense, *HBB, HBA2* and *HBA1* variants. Table 2 shows all threshold pairs that reach at least supporting level of evidence for both pathogenic and benign prediction in different SNV subsets. The full analysis for all thresholds and subsets is available in the Supplementary File 2 and the finetuning of the selected tools is available in Supplementary File 3.

**Figure 3.**
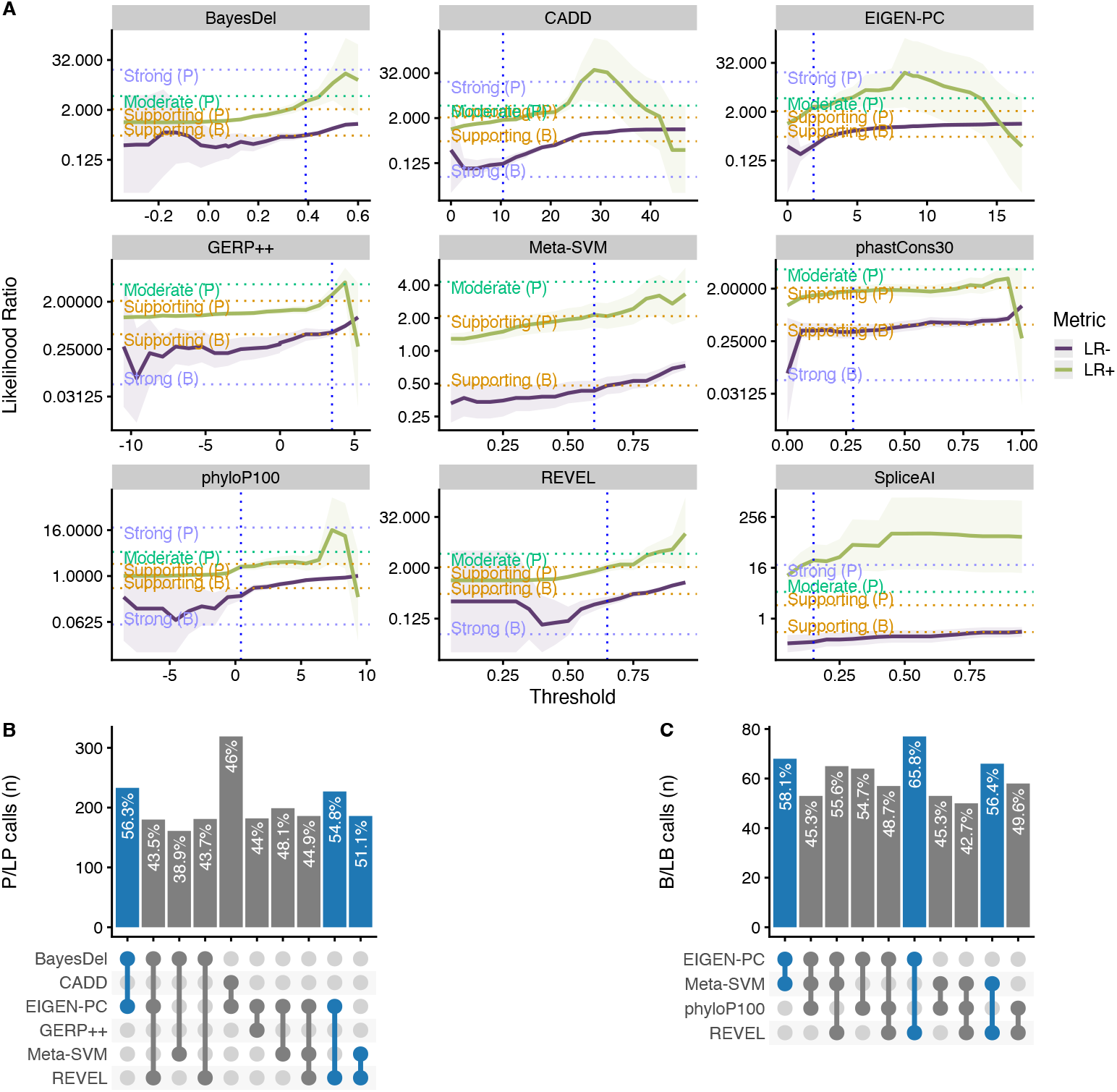
(A) Likelihood ratios of the top performing *in silico* tools with variable threshold. Vertical dashed lines indicate the optimal threshold based on the highest MCC. (B) Top ten tool combinations for pathogenic prediction based on concordance rate, with the top three shown in blue. (C) Top ten tool combinations for benign prediction based on concordance rate, with the top three shown in blue.

**Table 2.** *In sxilico* tools with pairs of non-overlapping thresholds that reach at least supporting evidence strength for both pathogenic and benign prediction. LR: likelihood ratio; Cl: confidence interval; PV: predicted variants

| Tool | Pathogenic threshold | LR+ | LR+ 95% CI | Strength (Pathogenic) | Benign threshold | LR- | LR- 95% CI | Strength (Benign) | Correctly PV | % of correctly PV |
| --- | --- | --- | --- | --- | --- | --- | --- | --- | --- | --- |
| All SNVs |  |  |  |  |  |  |  |  |  |  |
| BayesDel_addAF | ≥0.39 | 3.21 | [2.19, 4.72] | Supporting | <0.23 | 0.35 | [0.26, 0.47] | Supporting | 302 | 56.87 |
| CADD | >25 | 8.27 | [4.34, 15.75] | **Moderate** | ≤21.75 | 0.42 | [0.37, 0.48] | Supporting | 418 | 47.18 |
| CADD | >16.3 | 2.59 | [2.1, 3.2] | Supporting | ≤16.3 | 0.26 | [0.21, 0.31] | Supporting | **703** | **79.35** |
| Eigen-PC | >1.9 | 3 | [2.21, 4.07] | Supporting | ≤1.9 | 0.28 | [0.22, 0.35] | Supporting | 415 | 78.15 |
| GERP++ | >4.22 | 4.33 | [2.51, 7.49] | Supporting | ≤0.15 | 0.32 | [0.22, 0.46] | Supporting | 225 | 42.37 |
| MetaSVM | >0.81 | 3.25 | [2.16, 4.89] | Supporting | ≤0.46 | 0.38 | [0.3, 0.48] | Supporting | 272 | 56.55 |
| phyloP100way | >7.32 | 17.8 | [2.5, 127] | Supporting | ≤0.8 | 0.36 | [0.28, 0.46] | Supporting | 130 | 24.48 |
| REVEL | >0.77 | 3.05 | [2.12, 4.4] | Supporting | ≤0.7 | 0.38 | [0.31, 0.47] | Supporting | 309 | 64.24 |
| SpliceAI | >0.3 | 58.12 | [27.23, 124.03] | **Strong** | ≤0.3 | 0.33 | [0.23, 0.48] | Supporting | 637 | 96.08 |
| Missense only |  |  |  |  |  |  |  |  |  |  |
| BayesDel_addAF | ≥0.41 | 3.35 | [2.2, 5.12] | Supporting | <0.22 | 0.32 | [0.23, 0.45] | Supporting | 241 | 51.72 |
| CADD | >23.25 | 3.19 | [2.17, 4.69] | Supporting | ≤20.9 | 0.36 | [0.28, 0.46] | Supporting | 283 | 60.6 |
| Eigen-PC | >1.9 | 2.93 | [2.16, 3.98] | Supporting | ≤1.9 | 0.3 | [0.24, 0.38] | Supporting | **357** | **76.61** |
| GERP++ | >4.22 | 4.27 | [2.47, 7.4] | Supporting | ≤-0.87 | 0.31 | [0.2, 0.47] | Supporting | 187 | 40.13 |
| MetaSVM | >0.8 | 3.08 | [2.09, 4.53] | Supporting | ≤0.39 | 0.37 | [0.29, 0.48] | Supporting | 267 | 57.3 |
| phastCons30way | >0.94 | 3.19 | [2.09, 4.88] | Supporting | ≤0.41 | 0.36 | [0.28, 0.46] | Supporting | 252 | 54.08 |
| phyloP100way | >7.32 | 19.11 | [2.68, 136.47] | Supporting | ≤0.56 | 0.35 | [0.27, 0.46] | Supporting | 119 | 25.54 |
| REVEL | >0.77 | 3.02 | [2.09, 4.35] | Supporting | ≤0.7 | 0.39 | [0.32, 0.48] | Supporting | 297 | 63.73 |
| Non-missense only |  |  |  |  |  |  |  |  |  |  |
| CADD | >11.5 | 8.62 | [3.42, 21.77] | Supporting | ≤11.5 | 0.08 | [0.05, 0.11] | Supporting | 350 | 92.84 |
| SNVs in *HBB* |  |  |  |  |  |  |  |  |  |  |
| BayesDel_addAF | ≥0.31 | 6.43 | [2.23, 18.58] | Supporting | <0.31 | 0.22 | [0.17, 0.3] | Supporting | 210 | 81.08 |
| CADD | >25.25 | 31.64 | [4.5, 222.38] | **Moderate** | ≤22.65 | 0.42 | [0.37, 0.48] | Supporting | 264 | 48.71 |
| CADD | >10.8 | 3.26 | [2.29, 4.64] | Supporting | ≤10.8 | 0.08 | [0.05, 0.12] | Supporting | 494 | 91.14 |
| SNVs in *HBA1* |  |  |  |  |  |  |  |  |  |  |
| CADD | >22.95 | 4.94 | [2.29, 10.68] | Supporting | ≤17 | 0.3 | [0.19, 0.48] | Supporting | 84 | 61.76 |

Notably, CADD is the only tool that reached a moderate level of evidence (LR+ lower bound 95% CI ≥ 4.3) for prediction of pathogenic variants (threshold ≥ 25), while BayesDel, Eigen-PC, GERP++, REVEL, MetaSVM, phyloP100way and CADD (with a lower threshold of 16.3) have also reached the supporting evidence strength. Importantly, CADD (at Supporting strength), Eigen-PC and REVEL correctly predict the highest number of SNVs with 79.35%, 78.15% and 64.24%, respectively. Moreover, SpliceAI reached strong level of evidence for splicing prediction (threshold > 0.3), correctly predicting 96.08% of all variants.

When evaluating the performance of tools on the subset of missense variants, we identified eight tools (BayesDel, Eigen-PC, GERP++, MetaSVM, REVEL, CADD, phyloP100way and phastCons30way) that reached supporting strength level. Eigen-PC, REVEL, and CADD achieved the highest percentages of correctly predicted SNVs with 76.61%, 63.73%, and 60.6%, respectively. Moreover, CADD performed well for non-missense variants where a single threshold of 11.5 produced an accuracy of 92.84%, while achieving supporting strength.

With regards to the gene-specific analysis, BayesDel and CADD performed well for the prediction of *HBB* variants using a single threshold and accuracies of 81.08% and 91.14%, respectively, with CADD achieving moderate strength for pathogenic prediction with a threshold of 25.25. Furthermore, CADD achieved supporting strength for SNVs in *HBA1*, whilst no tool reached the required LR thresholds for *HBA2.*

Figures 2B and 2C show the concordance among the top performing tools of this study for pathogenic and benign prediction, respectively, using the recommended thresholds shown in Table 2 (full dataset; supporting strength thresholds). Although the overall concordance is low, some tools, such as Eigen-PC and REVEL, have higher concordance rates for both pathogenic (54.8%) and benign (65.8%) prediction. The low concordance rate of the top performing tools is also reflected in the prediction of VUS (Supplementary Figure 3), where differences in the distribution of predicted pathogenicity classes are observed among *in silico* tools. Nonetheless, this will be further assessed when the pathogenicity status of these SNVs is clarified.

## Discussion

The main goal of this study was to assess the performance of *in silico* prediction tools in the context of haemoglobinopathy-specific SNVs and to provide evidence to the ClinGen Hemoglobinopathy VCEP for the most appropriate use of computational evidence in variant interpretation based on the ACMG/AMP guidelines. We evaluated the performance of 31 *in silico* predictors on a set of 1627 haemoglobinopathy-specific SNVs. The pathogenicity of these variants was assessed using a Delphi approach by haemoglobinopathy experts based on literature review and experimental evidence.

Our comparative analysis showed that, when used as binary predictors of pathogenicity, most tools have high sensitivity and accuracy but suffer from poor specificity. We show that binary classification results in low LRs for most tools and, thus, cannot be used alone based on the Bayesian Framework for variant classification^54^. Instead, as we demonstrated in this study, stronger evidence is obtained when we trichotomized the problem by independently defining different non-overlapping thresholds for pathogenic and benign prediction of globin gene variants. This approach was previously described by other ClinGen VCEPs, evaluating sequence variants in other genes^56^. Our findings show that Eigen-PC, REVEL and CADD performed well for predicting the functional effect of missense SNVs, while CADD was also a strong predictor of non-missense variants. Meta-predictors BayesDel and MetaSVM were also strong performers in our comparison, while GERP++, phyloP100way, and phastCons30way performed better among the conservation tools, albeit with a lower overall accuracy. Out of the four splicing prediction tools evaluated, SpliceAI performed better and produced the highest LR+ values reaching strong level of evidence. However, due to the low number of negative examples in our dataset for the other splicing tools evaluated, these results should be interpreted with caution. Our results show that SpliceAI is a reliable predictor of the splicing impact of SNVs in the globin genes.

The annotated pathogenicity of the variants in our dataset was based on criteria agreed by all coauthors of this paper. These criteria are not based on the ACMG/AMP framework, because there is currently no available standard for pathogenicity classification of globin gene variants. The ClinGen Hemoglobinopathy VCEP is currently piloting its ACMG/AMP specifications, which can be used for variant classification in the future, thus potentially leading to reassessment of *in silico* predictors for globin genes variants. Nevertheless, the current classification reflects the current knowledge about the pathogenicity of the variants in our dataset, agreed by experts involved in the molecular diagnosis of haemoglobinopathies in five countries (Cyprus, Greece, Malaysia, Netherlands, and Portugal). A potential limitation is that some benign variants have not been observed *in trans* with both a β-thalassaemia variant and the Hb S variant and, therefore, their pathogenicity is assigned based on the current knowledge in the field. However, our approach is justified, because small numbers of true benign SNVs reflect the reality in clinical diagnostics, where pathogenic SNVs associated with clinical phenotypes- are more easily interpreted than benign ones.

This study provides evidence for the selection of the most suitable *in silico* tools for the interpretation of SNVs in the globin gene clusters using the ACMG/AMP guidelines. Specifically, we provide the optimal thresholds for different tools that can be used under the PP3/BP4 criteria, including missense and splicing variant interpretation, while optimal thresholds for conservation-based tools are also critical for the application of criterion BP7. To our knowledge, this is the largest study evaluating the disease-specific application of *in silico* predictors in variant classification under the ACMG/AMP framework and its associated

Bayesian framework. Our approach can be further expanded for the optimal calibration of thresholds of *in silico* tools in other genes and diseases, hence facilitating variant interpretation using the ACMG/AMP framework.

## Supplementary Material

**Supplementary File 1.** The list of ClinGen Hemoglobinopathy VCEP members **Supplementary File 2.** Table with the dataset used in this study and the resulting scores obtained by the *in silico* predictors, divided into different sheets and subsets: all SNVs, missense only, non-missense only, *HBB, HBA1,* and *HBA2.*

**Supplementary File 3.** Refined thresholds for the nine selected *in silico* predictors, divided into different subsets: all SNVs, missense only, non-missense only, *HBB, HBA1,* and *HBA2*. Only decision thresholds passing the LR criteria are shown.

**Supplementary Figure 1.** Distribution of variants on each globin gene based on their actual pathogenicity. A bin size of 3 bp (inframe) and 5 bp in exonic and intronic regions, respectively, is used for the illustration.

**Supplementary Figure 2.** Heatmap illustrating the concordance and clustering of *in silico* tools with respect to the variant type and globin gene.

**Supplementary Figure 3.** Prediction of VUS using thresholds for the full dataset (at Supporting strength)

## Data availability statement

All data used and generated in this study are available in the supplementary material.

## Acknowledgments

We thank the Cyprus Institute of Neurology and Genetics for computer equipment and for hosting ITHANET. The list of ClinGen Hemoglobinopathy VCEP members is provided in Supplementary File 1.

## Author Contributions

ST, MX, PK, and MK conceived and designed the study; CS and AM collected and curated data; CLH, CB, EF, JTS, NMY, FSA, EE, HHF, BAZ, and MK provided SNVs and performed the expert pathogenicity classifications; ST, MX, and ACK performed bioinformatics analyses; ST, MX and AM prepared figures; ST, MX and PK wrote the manuscript; all authors have read and approved the final version of the manuscript.

## Funding Information

This work was co-funded by the European Regional Development Fund and the Republic of Cyprus through the Research and Innovation Foundation (Project: EXCELLENCE/1216/256).

## Conflict of Interest Statement

The authors declare no conflicts of interest

## Notes

### Competing Interest Statement

The authors have declared no competing interest.

