## Supplementary figures and images for "Evaluation of *in silico* predictors on short nucleotide variants in *HBA1, HBA2* and *HBB* associated with haemoglobinopathies"

### Supplementary Figure 1

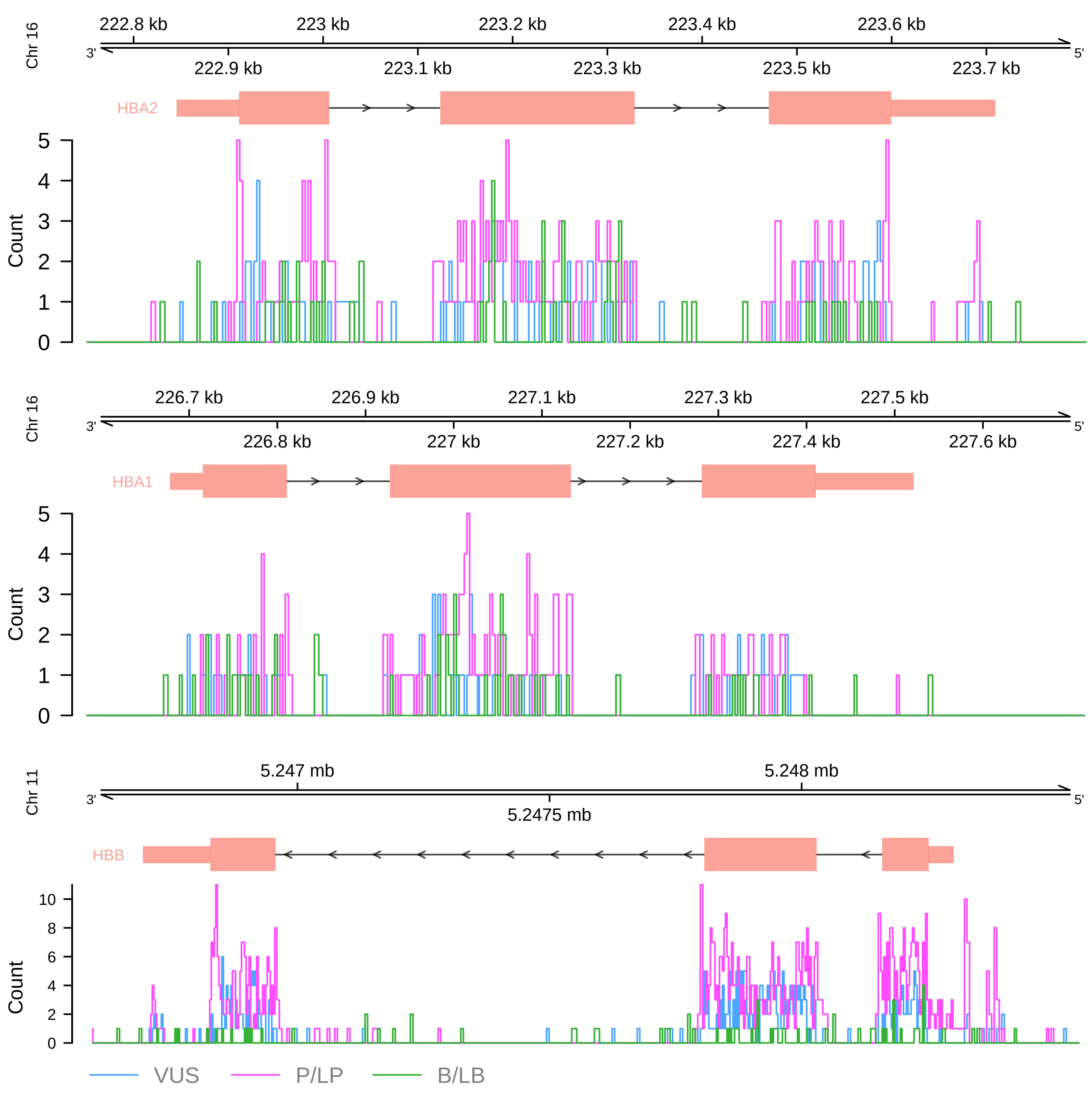

### Supplementary Figure 2

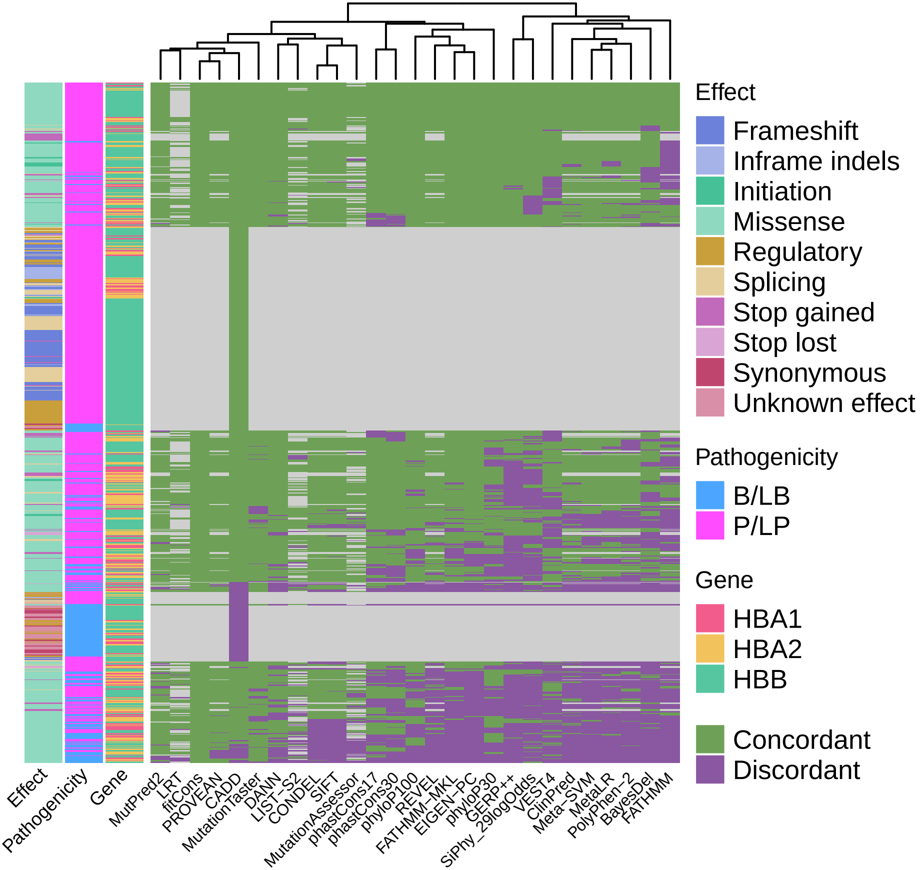

### Supplementary Figure 3

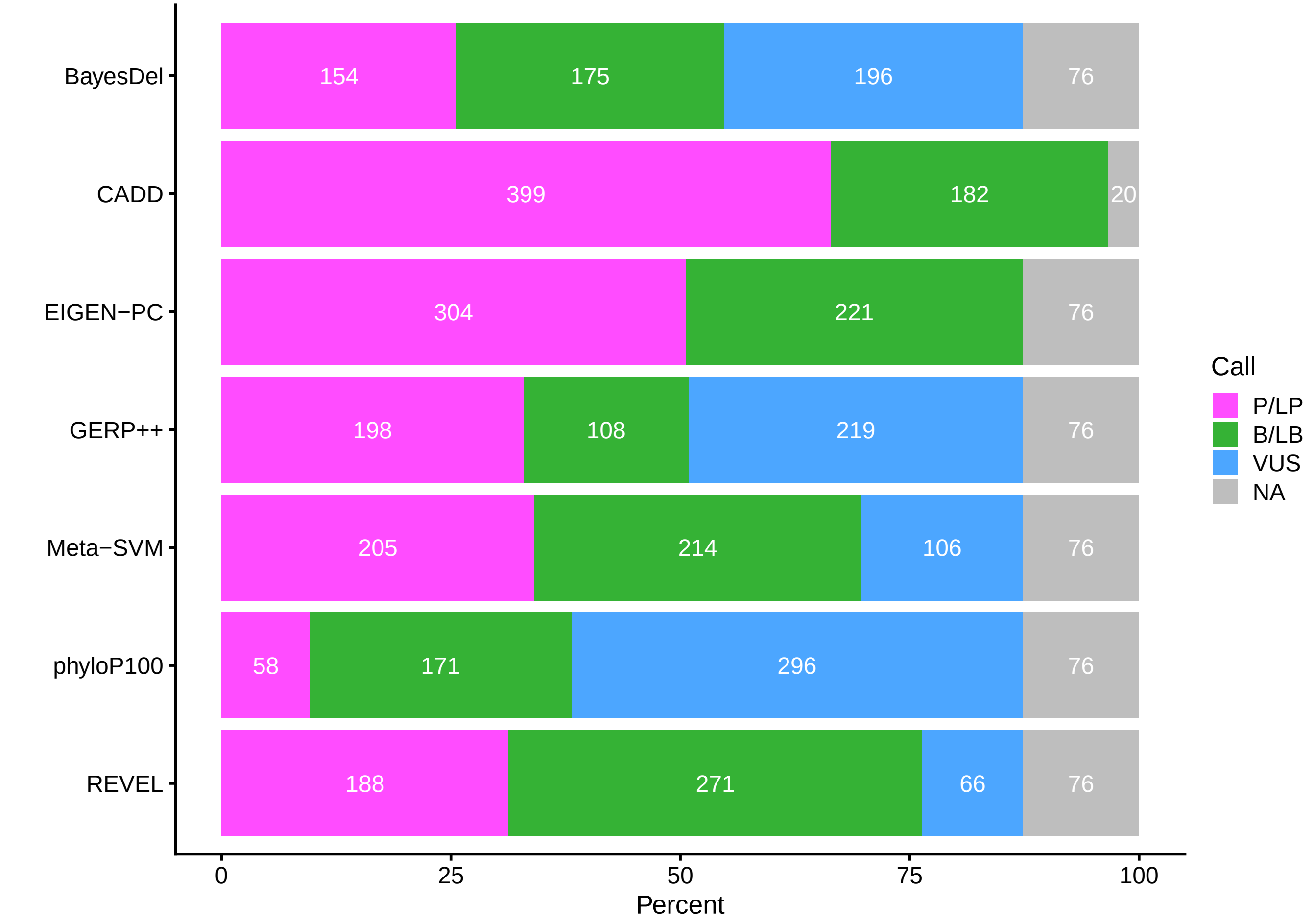
